# Task-irrelevant visual forms facilitate covert and overt spatial selection

**DOI:** 10.1101/2020.06.27.170894

**Authors:** Amarender R. Bogadhi, Antimo Buonocore, Ziad M. Hafed

## Abstract

Covert and overt spatial selection behaviors are guided by both visual saliency maps derived from early visual features as well as priority maps reflecting high-level cognitive factors. However, whether mid-level perceptual processes associated with visual form recognition contribute to covert and overt spatial selection behaviors remains unclear. We hypothesized that if peripheral visual forms contribute to spatial selection behaviors, then they should do so even when the visual forms are task-irrelevant. We tested this hypothesis in male and female human subjects as well as in male macaque monkeys performing a visual detection task. In this task, subjects reported the detection of a supra-threshold target spot presented on top of one of two peripheral images, and they did so with either a speeded manual button press (humans) or a speeded saccadic eye movement response (humans and monkeys). Crucially, the two images, one with a visual form and the other with a partially phase-scrambled visual form, were completely irrelevant to the task. In both manual (covert) and oculomotor (overt) response modalities, and in both humans and monkeys, response times were faster when the target was congruent with a visual form than when it was incongruent. Importantly, incongruent targets were associated with almost all errors, suggesting that forms automatically captured selection behaviors. These findings demonstrate that mid-level perceptual processes associated with visual form recognition contribute to covert and overt spatial selection. This indicates that neural circuits associated with target selection, such as the superior colliculus, may have privileged access to visual form information.

**Significance statement:** Spatial selection of visual information either with (overt) or without (covert) foveating eye movements is critical to primate behavior. However, it is still not clear whether spatial maps in sensorimotor regions known to guide overt and covert spatial selection are influenced by peripheral visual forms. We probed the ability of humans and monkeys to perform overt and covert target selection in the presence of spatially congruent or incongruent visual forms. Even when completely task-irrelevant, images of visual objects had a dramatic effect on target selection, acting much like spatial cues used in spatial attention tasks. Our results demonstrate that traditional brain circuits for orienting behaviors, such as the superior colliculus, likely have privileged access to visual object representations.

## Introduction

Spatial selection of stimuli in a cluttered visual scene is central to visual behaviors in primates, and it could occur either overtly (with orienting eye movements) or covertly (without such eye movements). The mechanisms underlying both overt and covert spatial selection behaviors rely on spatial maps in sensorimotor regions that are functionally organized into visual saliency maps, primarily derived from low-level visual processes, and priority maps, representing high-level cognitive processes (Fecteau and Munoz, 2006; Veale et al., 2017; Bisley and Mirpour, 2019). Indeed, classic visual saliency map models are computed from early visual features such as orientation, motion, and color (Itti and Koch, 2000), whereas, priority maps are based on cognitive factors such as behavioral relevance, expectation and reward (Awh et al., 2012; Chelazzi et al., 2014; Sprague et al., 2018). Accordingly, visual saliency maps and priority maps are believed to be represented in the neuronal activity of visual, sensorimotor, and associative brain regions, such as primary visual cortex (V1), superior colliculus (SC), and regions of the parietal (LIP) and prefrontal (FEF) cortices (Gottlieb et al., 1998; Bisley and Goldberg, 2003; Ignashchenkova et al., 2003; White et al., 2017; Sapountzis et al., 2018; Yan et al., 2018; Chen et al., 2020).

The organization of spatial maps based on a dichotomy of early visual features, on the one hand, and high-level cognitive factors, on the other, ignores whether mid-level perceptual processes related to visual form recognition are represented in these maps. This is inconsistent with multiple lines of evidence suggesting a potential link between visual form recognition and spatial selection behaviors. First, recent studies in monkeys identified a novel attention-related region in the temporal cortex (Bogadhi et al., 2019a; Stemmann and Freiwald, 2019). Importantly, neurons in this attention-related region were selective for peripheral visual forms (Bogadhi et al., 2019b), suggesting a functional link between covert spatial selection and peripheral visual form recognition. Second, studies modeling fixation patterns in free-viewing of natural images show that visual objects predict fixation patterns better than saliency maps based on early visual features (Einhäuser et al., 2008; Yanulevskaya et al., 2013; Kümmerer et al., 2014), indicating an influence of visual forms on overt behaviors. Third, behavioral studies in humans show a rapid detection of faces and animals in peripheral images for saccadic eye movements and attentional capture, suggesting a rapid processing of animate visual forms for overt selection (Kirchner and Thorpe, 2006; Bindemann et al., 2007; Crouzet et al., 2010; Drewes et al., 2011; Devue et al., 2012). However, such rapid detection could be explained by low-level image features or unnatural statistics of the image databases (Honey et al., 2008; Wichmann et al., 2010; Crouzet and Thorpe, 2011; Zhu et al., 2013). Importantly, these studies used visual form images as saccade targets, which were always relevant to the task performance. Hence, it remains unclear, from studies using goal-directed and free-viewing paradigms, if peripheral visual forms contribute to spatial selection when they are rendered irrelevant to the task and equated with non-form images for low-level visual features. We hypothesized that if peripheral visual forms contribute to spatial selection, then they should do so in both covert and overt behaviors, even when the visual forms are task-irrelevant and equated for low-level visual features.

We investigated the contribution of peripheral visual forms to covert and overt spatial selection using a visual detection task pitting visual form images against 50% phase-scrambled images. Most importantly, all images were irrelevant to the task and matched for low-level image properties. In the covert (humans) and overt (humans and monkeys) tasks, subjects reported the detection of a supra-threshold target with a manual or saccadic eye movement response, respectively. We found that response times were significantly faster when the target was congruent with a visual form image in both covert and overt selection tasks, and in both humans and monkeys. Crucially, almost all response errors were captured by visual forms incongruent with targets. Interestingly, during covert selection, microsaccades following image onsets were biased towards visual forms. These findings demonstrate that peripheral visual forms, even when task-irrelevant, contribute to overt and covert spatial selection and perhaps act as spatial cues for orienting movements (Posner, 1980; Tian et al., 2016).

## Materials and Methods

### Subjects and ethics approvals

11 human subjects (3 males and 8 females; mean age ± s.d. = 27.3 ± 3.9 years) naïve to the purpose of the study and 3 male rhesus monkeys (*Macaca mulatta*; monkeys A, F, and M) aged 10, 11, and 10 years, respectively, participated in this study. All human subjects provided written informed consent in accordance with the Declaration of Helsinki. Ethics committees at the Medical Faculty of Tuebingen University reviewed and approved protocols for the human experiments. Monkey experiments were approved by regional governmental offices in Tuebingen.

### Experimental setups

Human subjects were seated in a dark room at a viewing distance of 57 cm from a cathode-ray tube (CRT) monitor with a resolution of 1400 x 1050 pixels (34.13° x 25.93°). Stimulus display on the monitor was controlled by a 2010 Mac Pro (Apple Inc., Cupertino, CA) running MATLAB (The Mathworks, Natick, MA) with the Psychophysics Toolbox extensions (Brainard, 1997). Eye position signals and manual responses were acquired using an EyeLink 1000 infrared eye-tracking system (SR Research Ltd., Ottowa, Ontario, Canada) and Viewpixx button box (VPixx Technologies, Saint-Bruno, QC Canada), respectively.

Monkeys were seated and head-fixed in a primate chair (Crist Instrument Inc., Hagerstown, MD) inside a darkened booth at a viewing distance of 72.2 cm from a CRT monitor with a 1024 x 768 resolution (30.96° x 23.47°). For experiments in monkeys A and M, stimulus display on the monitor was controlled using a modified version of PLDAPS with Datapixx and Psychophysics Toolbox extensions on MATLAB (The Mathworks, Natick, MA) running on an Ubuntu operating system (Eastman and Huk, 2012). For experiments in monkey F, stimulus display was controlled using a LabVIEW system (National Instruments, Austin, TX) handshaking with a 2010 Mac Pro (Apple Inc., Cupertino, CA) running MATLAB (The Mathworks, Natick, MA) with the Psychophysics Toolbox extensions (see Chen and Hafed, 2013; Tian et al., 2016 for details). Eye position signals in monkeys A and M were measured using a surgically implanted scleral search coil; eye position signals in monkey F were measured using an EyeLink 1000 infrared eye-tracking system (SR Research Ltd., Ottowa, Ontario, Canada). Surgical procedures for implanting head-holders and scleral coils were described in a previous study (Skinner et al., 2019).

### Experimental design: covert selection task (human subjects)

Subjects started each trial by fixating a central spot of 0.1° radius (97.6 cd/m^2^) displayed on a gray background (43.84 cd/m^2^). Eye position signals were monitored to enforce fixation within a fixation window of 2° radius. Following a 500 – 1000ms randomized fixation duration, an intact visual form image (4.88° x 4.88°) and its corresponding 50% scrambled form image (see *Image normalization* below) were displayed symmetrically on either side of fixation along the horizontal meridian and centered at 8° eccentricity. After a fixed delay of 100ms, 200ms, or 300ms following image onset, a supra-threshold target (black disc; radius = 0.2°) was displayed at the center of one of the two images. Subjects were instructed to report the spatial location of the target with a left or right button press, and most importantly, they were informed that both images were irrelevant to the detection task. We refer to trials in which the target was presented on top of the visual form image as target congruent trials and trials in which the target was presented on top of the 50% phase-scrambled image as target incongruent trials. All three delay conditions (100ms, 200ms, or 300ms) were randomized across trials. In addition, catch trials with no target were also included in the 100ms and 300ms delay conditions on 25% of trials, and subjects were instructed to withhold their responses in these trials. Data in each subject (n = 11) were collected in a single session.

### Experimental design: overt selection task (humans and monkeys)

The trial structure in the overt task was the same as in the covert task with one difference: in the overt task, subjects reported the detection of the target with a saccade to the target location rather than a button press. We also included a single image condition on 40% of trials. In this case, only one image, either a visual form image or a 50% scrambled form image, was presented simultaneously with a target on top of it in one of the four diagonal locations at 8° eccentricity. The single image condition with diagonal locations was used to control for any spatial biases in eye movements from repetitive target presentations at the same spatial locations. We collected data in 10 of the 11 human subjects in a single session each and in 3 monkeys across 17 sessions. In monkeys, we used the same visual images (now sized 5.14° x 5.14°) as in the human experiments, with a target (black disc) of radius 0.3° - 0.45° across monkeys. The background gray luminance for the monkey experiments was 27.21 – 37.1 cd/m^2^. The humans performed the overt selection task before performing the covert selection task.

### Image normalization

Forty images with visual forms and their corresponding 50% scrambled form images were used in both human and monkey experiments. Visual form images were obtained from previous electrophysiology studies of the inferotemporal cortex (Tsao et al., 2006; Bogadhi et al., 2019b) and were sampled from four different categories including human faces, fruits, bodies, and inanimate objects with ten examples in each category (see Fig. 1c in Results).

**Figure 1.**
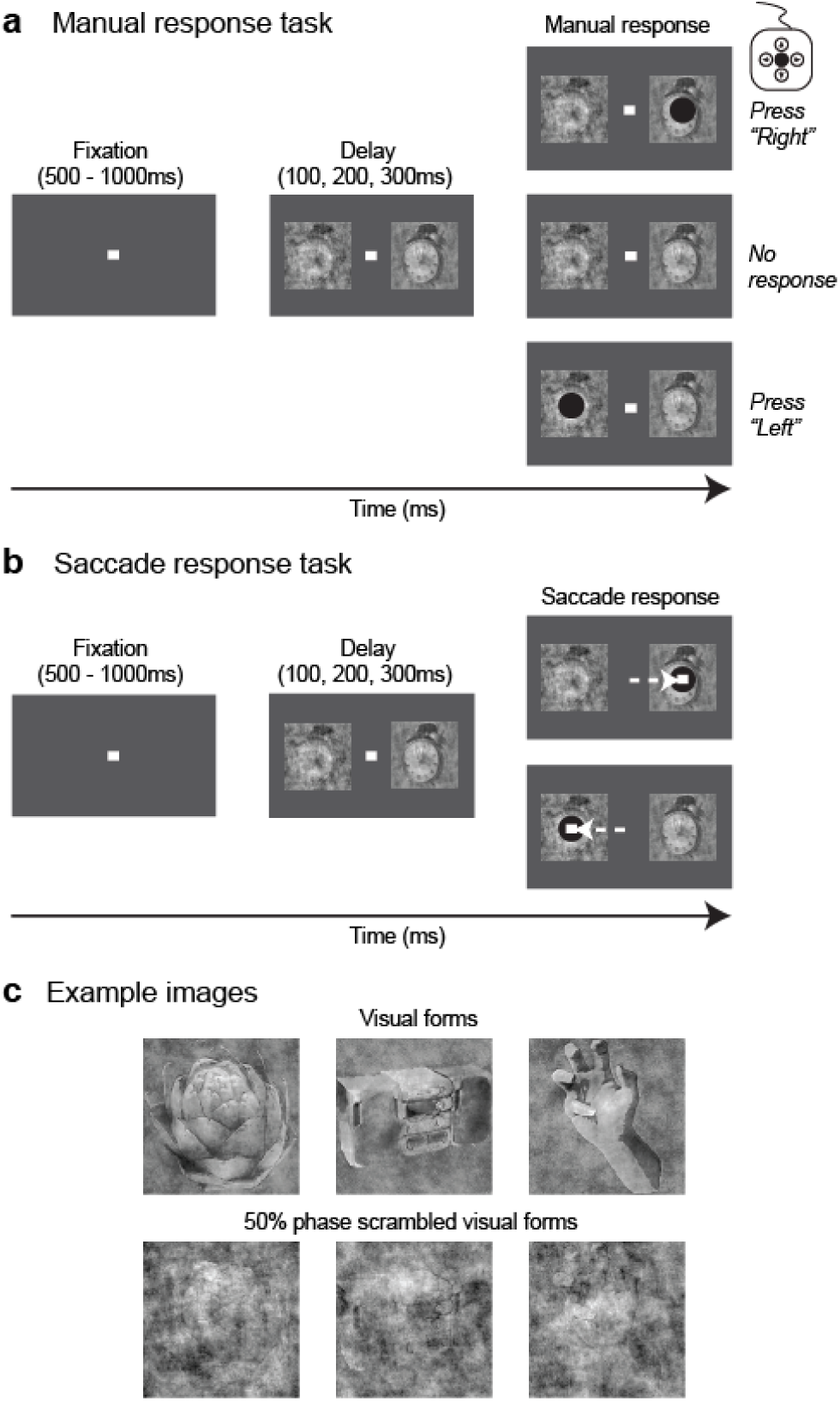
Covert (manual response) and overt (saccade response) selection tasks. **(a)** Trial epochs in the covert selection task. Each trial started with a fixation spot at the center of the screen. After 500 – 1000ms of fixation, two images, one with an intact visual form and the other with the corresponding 50% phase-scrambled visual form, appeared on either side of fixation along the horizontal meridian and centered at 8° eccentricity. Following a fixed delay of either 100ms, 200ms, or 300ms, a supra-threshold target (“black disc”) appeared on top of one of the images and subjects were instructed to report the detection of the target with an appropriate button press. On the remaining “catch” trials (see Methods), no target was presented, and subjects were required to withhold their response. **(b)** Trial epochs in the overt selection task. The task consisted of instruction (“single image”) and distractor (“two images”) trials. The distractor trials shown here were similar to the covert selection task until the target presentation. After a fixed delay of 100ms, 200ms, or 300ms from target onset, a supra-threshold target (“black disc”) appeared on top of one of the images, and the subjects were instructed to detect the target and generate a saccade to its location. At the time of target appearance, the fixation spot at the center was extinguished and was presented on top of the target to instruct the subjects to generate a visually-guided saccade. **(c)** Example images of intact visual forms (top row) and their corresponding 50% phase-scrambled visual forms (bottom row) from 3 of the 4 categories used in this study. An example image from the fourth “face” category is not shown here. A total of 40 visual form images, 10 from each category, were used after equating for low-level features (see Methods).

All images were equated iteratively for luminance distributions (mean = background luminance) and Fourier spectra using the SHINE tool box (Willenbockel et al., 2010). Briefly, all forty visual form images were resized to the appropriate dimensions, and the mean gray level of each image was equated to the background level (*lumMatch in SHINE*). The resultant forty images were iteratively (n = 20) matched for the histogram of gray levels (*histMatch in SHINE*) and the Fourier spectra (*specMatch in SHINE*), before generating their corresponding 50% phase-scrambled images by randomizing half of the phase matrix and keeping the amplitude matrix constant. Finally, all of the visual form images and their corresponding phase-scrambled images were iteratively (n = 20) matched, once again, for the histogram of gray levels and the Fourier spectra to yield the final visual form images (see top panel in Fig. 1c) and phase-scrambled images (see bottom panel in Fig. 1c) used in this study.

### Supra-threshold target detection

We hypothesized that visual form contribution to spatial selection behaviors, in covert and overt tasks, should be evident as a facilitation of response times between target congruent and target incongruent conditions. Hence, it was important that the differences in response times between target congruent and target incongruent conditions could not be attributed to difficulty in the visual detection of the target across conditions. For this reason, we chose a high contrast (“black” color) and sufficiently large target (0.2° radius disc). We also confirmed that detection performance during the most difficult condition of our covert task (the 100ms delay condition; see Results) was supra-threshold in both target congruent (% correct performance = 99.48% ± 0.97% s.d.) and target incongruent (% correct performance = 99.06% ± 0.94% s.d.) trials, with no significant difference between the two conditions (Wilcoxon signed-rank test, p = 0.43).

### Statistical analyses: response times and proportions of errors

We measured visual form contribution to spatial selection behaviors on correct and error trials separately. Correct trials were defined as the trials in which the first response of the subject correctly matched the target location. Error trials were those in which the subjects erroneously selected the image that had no target dot superimposed on it (i.e. they selected the image opposite to the target). On correct trials, we quantified the effect of visual forms on response time differences between target congruent and target incongruent conditions. On error trials, we quantified the effect of visual forms on the proportion of errors between target congruent and target incongruent conditions. Response time on a correct trial was calculated as the time of response onset relative to target onset. The proportion of errors in each subject was calculated as the ratio of the number of error trials to the sum of correct and error trials.

Saccadic responses to targets in the overt task and microsaccades during fixation in both the covert and overt tasks were detected using a velocity and acceleration threshold followed by manual inspection (Krauzlis and Miles, 1996). Trials with microsaccades occurring between 100ms before image onset and response onset were excluded in the analyses of response times and proportion of errors to control for the effects of microsaccades on stimulus onset activity in different brain regions (Chen et al., 2015).

For analysis of response times in the covert task in humans (e.g. see Fig. 2a in Results), we included an average of 80.6 (s.d. = 26.4) and 77.9 (s.d. = 24.9) trials from target congruent and target incongruent conditions, respectively, for a given delay and subject. Similarly, in the overt task (Fig. 2b in Results), we included an average of 68.2 (s.d. = 9.8) and 68.6 (s.d. = 9.8) trials in target congruent and target incongruent conditions, respectively. For the response time analysis in monkeys, trial counts in target congruent and target incongruent conditions are shown in Fig. 6 in Results. For all paired comparisons in humans and monkeys, we used Wilcoxon signed-rank tests.

**Figure 2.**
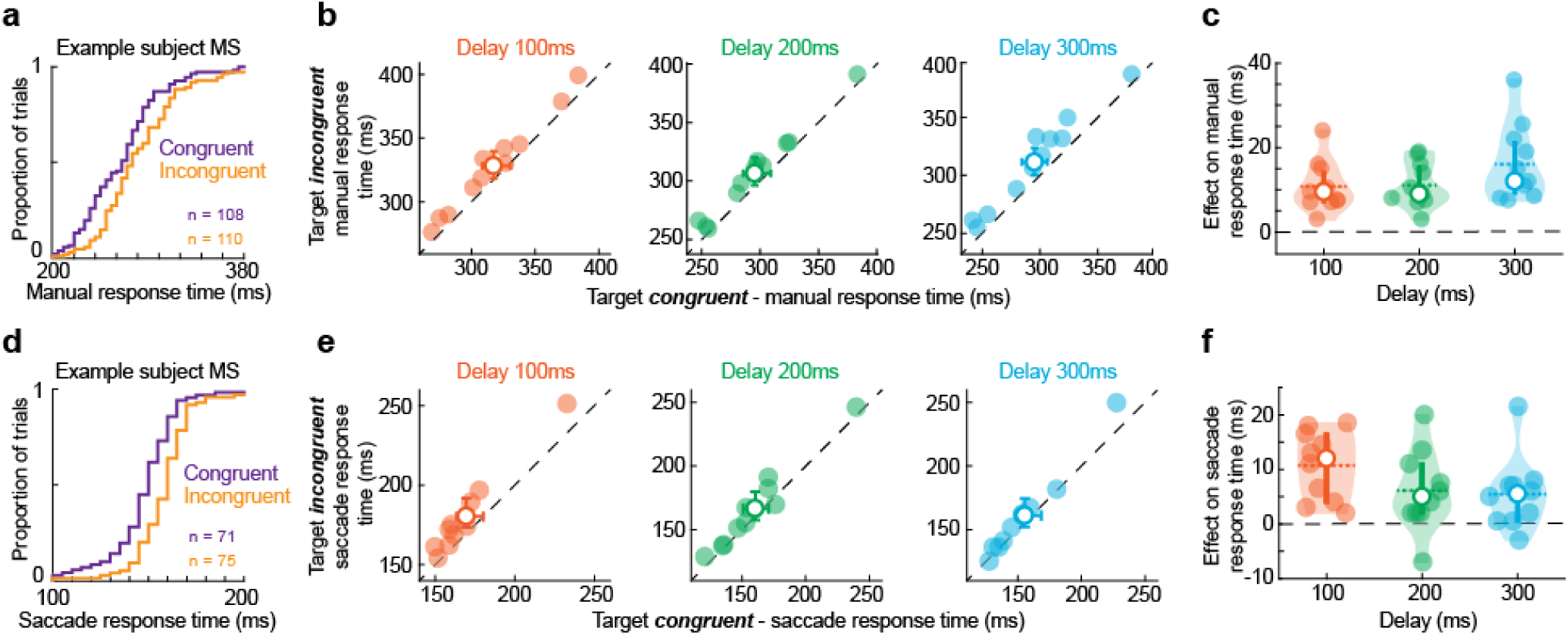
Visual form facilitation of response times in covert and overt selection tasks in humans. **(a)** Cumulative distribution of manual response times in an example subject for target congruent and target incongruent trials in the covert selection task during the fixed delay interval of 100ms (see Methods). The number of samples in each cumulative distribution is shown in the panel. **(b)** Each panel shows a paired comparison of median response times between target congruent and target incongruent trials in the covert selection task across all subjects (n = 11; filled circles) for a fixed delay of 100ms (red), 200ms (green), or 300ms (blue). The colored circle with errors bars in each panel represents mean and standard deviation across subjects. Dotted line indicates the line of unity slope; data above the line indicated faster response times in the target congruent condition. **(c)** The facilitatory effect of visual forms on manual response times in the covert task was quantified in each subject as a difference in median response times between target incongruent and target congruent trials (same data as in **b**). Each data point in the violin plots is from each subject. Colored circles and horizontal bars in each violin plot indicate median and mean, respectively. Vertical bars indicate standard deviation. Same color conventions as in **b. (d-f)** Same as in **a**-**c**, but now for the saccade responses of humans in the overt selection task.

**Figure 3.**
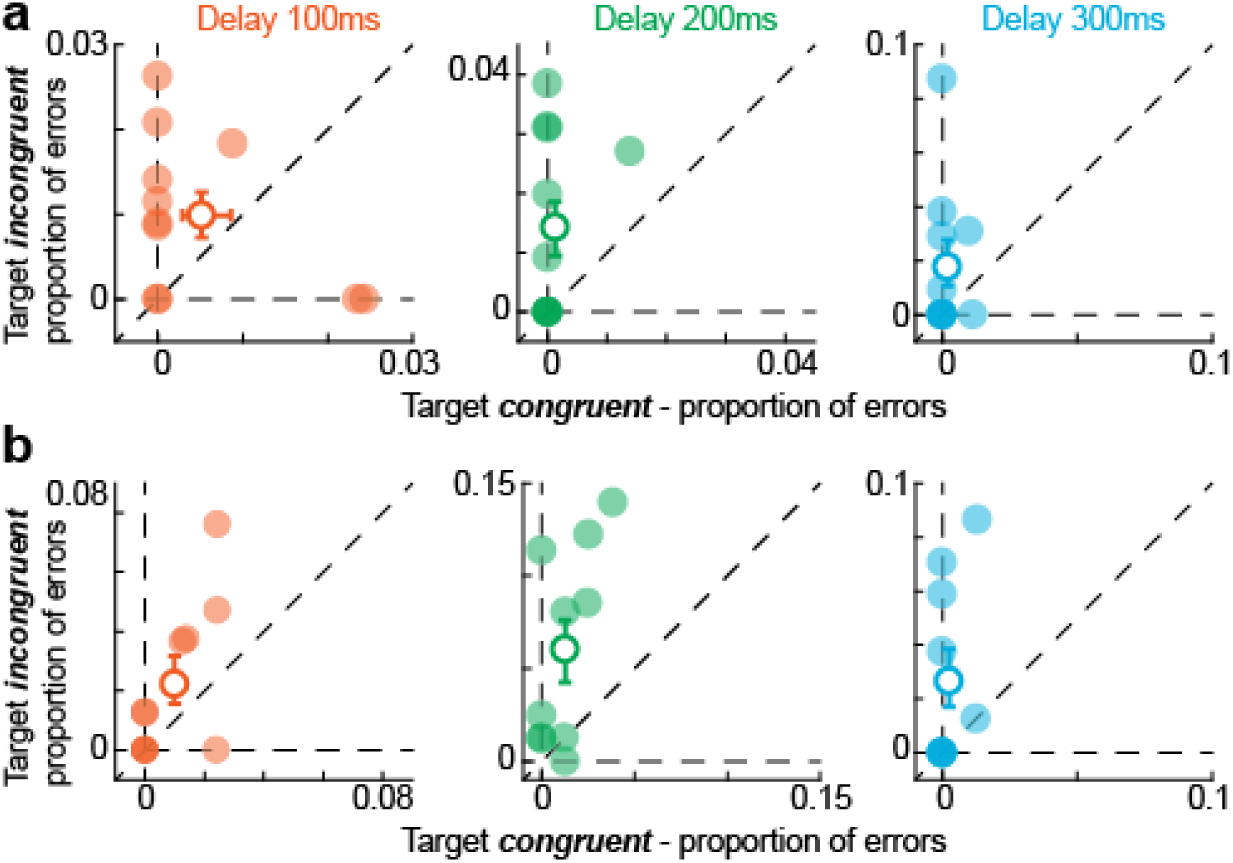
Visual form influence on errors in the covert and overt selection tasks in humans. **(a)** Each panel shows a paired comparison of the proportion of manual response errors between target congruent and target incongruent trials in the covert selection task across all subjects (n = 11; filled circles) for a fixed delay of 100ms (red), 200ms (green), or 300ms (blue). The colored circle with errors bars in each panel represents mean and standard deviation across subjects. Dotted line indicates line of unity slope; data above the line indicate more errors in the target incongruent condition. **(b)** Same as **a**, but now for the saccade response errors in the overt selection task.

**Figure 4.**
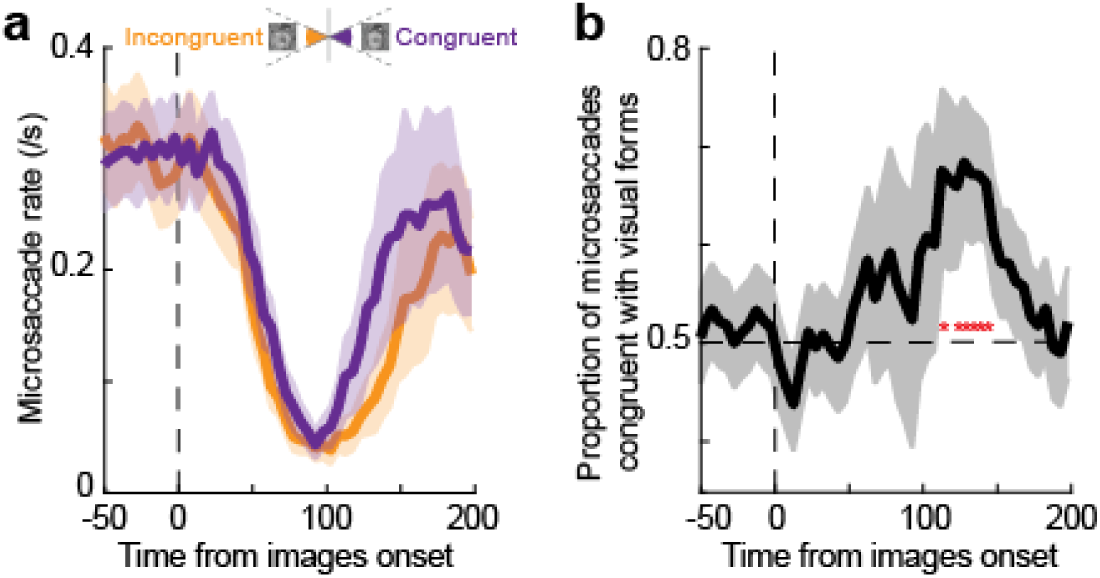
Visual form influence on human microsaccade directions in the covert selection task. **(a)** Mean microsaccade rate aligned to the time of image onset (vertical dotted line). Each curve shows the rate of microsaccades that were in the same (congruent; orange) or opposite (incongruent; purple) directions from the visual form image (see Methods). The inset shows a cartoon of visual angle (32°) subtended by the images for calculating congruent and incongruent microsaccade directions. Error bars indicate boot-strapped 68.2% confidence intervals. **(b)** Microsaccades in the congruent and incongruent directions (same data as in **a**) were used to show the proportion of microsaccades biased in the direction of visual forms 100ms after image onset (see Methods). Horizontal dotted line indicates chance-level. Error bars indicate boot-strapped 68.2% confidence intervals. Red colored symbols (*) indicate significance in the corresponding time bin using the binomial test (p < 0.05; see Methods).

**Figure 5.**
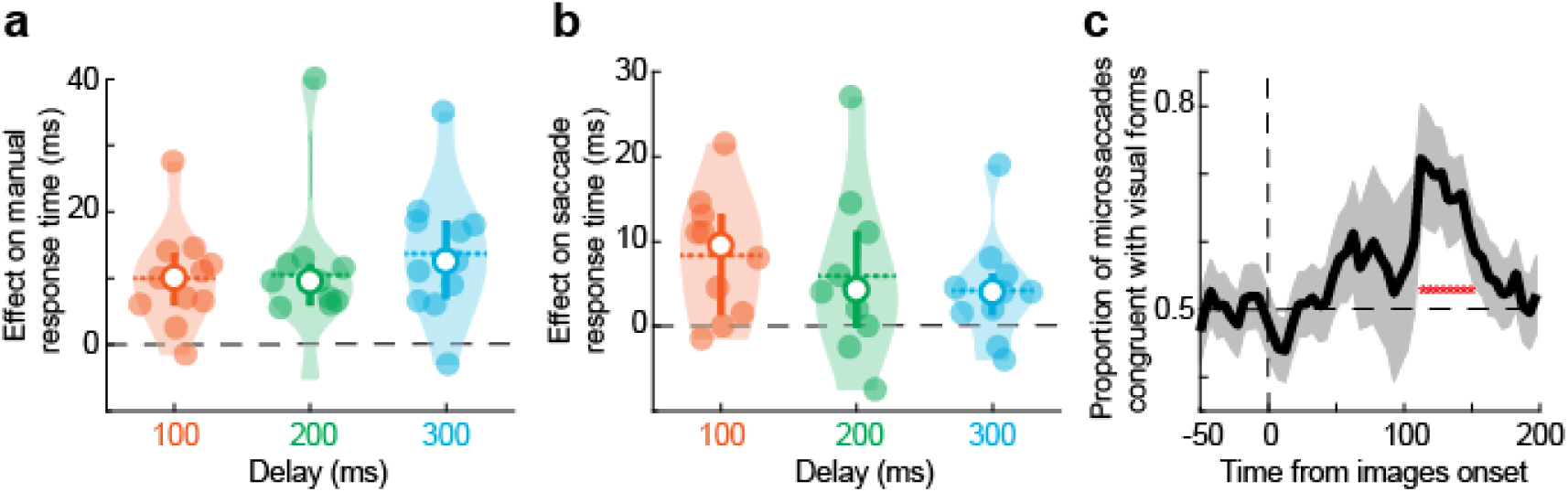
Non-face visual form facilitation of response times in covert and overt selection tasks in humans. **(a)** A facilitatory effect of non-face visual forms on manual response times in the covert selection task for the three delay conditions. Same conventions as in Fig. 2c. **(b)** A facilitatory effect of non-face visual forms on saccade response times in the overt selection task for the three delay conditions. Same conventions as in Fig. 2f. **(c)** Proportion of microsaccades biased in the direction of non-face visual forms after image onset, but before target onset (see Methods). Error bars indicate boot-strapped 68.2% confidence intervals. Red colored symbols in **c** (*) indicate significance in the corresponding time bin using the binomial test (p < 0.05; see Methods). Same conventions as in Fig. 4b.

**Figure 6.**
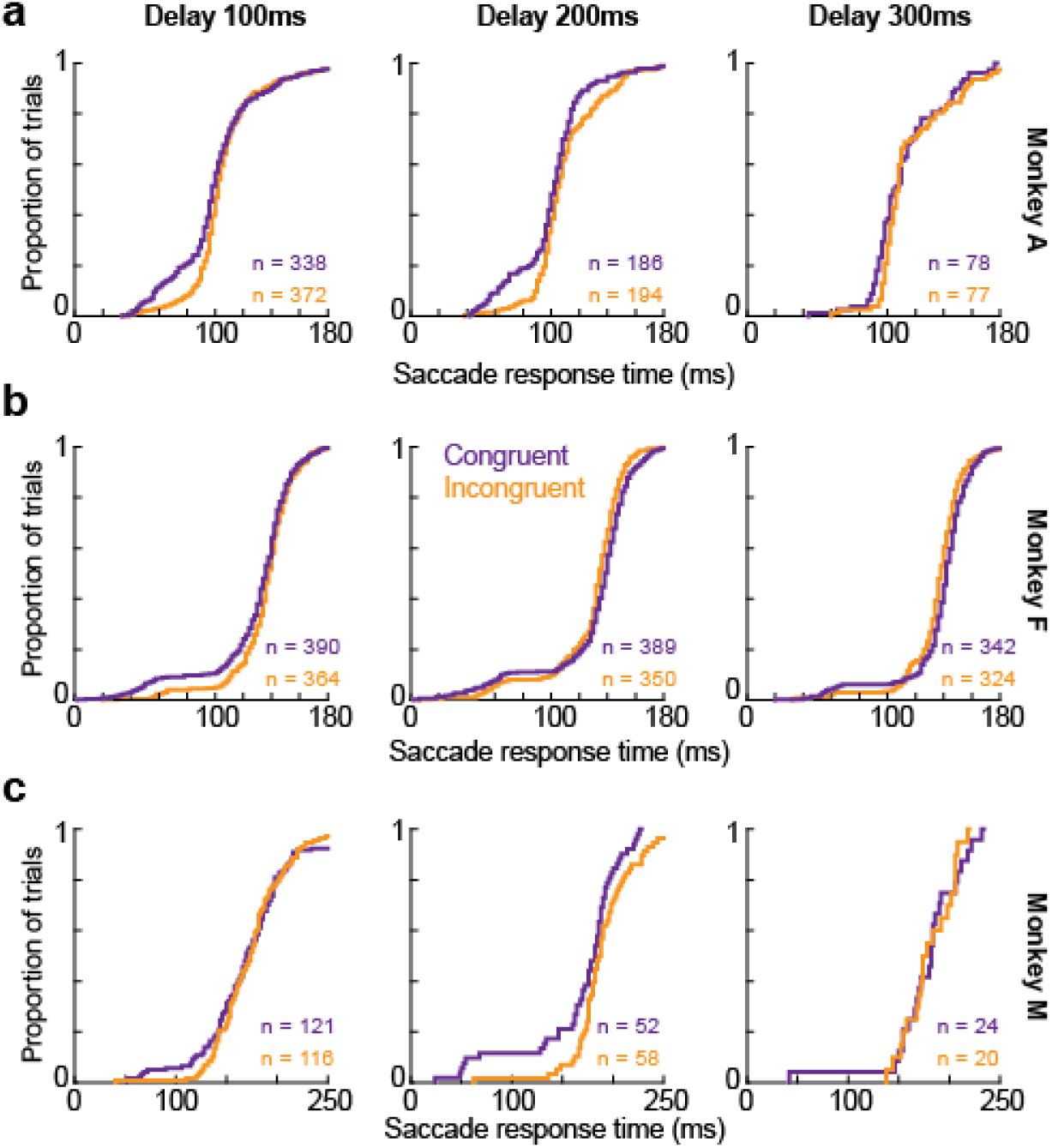
Distributions of saccade response times in the overt selection task in monkeys. **(a-c)** Cumulative distributions of saccade response times for target congruent and target incongruent trials in the overt selection task, similar to Fig. 2a, d in the example human subject. Each column represents a given delay condition (100ms: left, 200ms: middle, and 300ms: right columns), and each row represents a monkey (A, F, and M, respectively). The number of samples in each cumulative distribution is shown in the corresponding panel, and the color legend in the middle of the figure applies to all panels. All monkeys showed earlier response times for target congruent trials than for target incongruent trials when the response times were fast, but not when either the response times or target delay (300ms) were long. This is a magnified effect of the human observations in Fig. 2f.

### Statistical analyses: microsaccades

We pooled microsaccades flagged during the covert task across subjects, and we separated them into congruent and incongruent microsaccades based on their directions relative to the visual form image or the scrambled image, respectively. Specifically, we found that the great majority of microsaccades were predominantly horizontal due to our image configuration in the task. We therefore grouped all microsaccades within ±32deg from horizontal into two groups depending on the direction of their horizontal component: either towards the visual form image (congruent microsaccades) or towards the scrambled image (incongruent microsaccades; see inset in Fig. 4a in Results). The rate of congruent and incongruent microsaccades was constructed by counting the corresponding microsaccades in a 50ms time window sliding in steps of 5ms. The proportion of microsaccades congruent with visual forms was also calculated as the ratio of the number of congruent microsaccades to the total sum of congruent and incongruent microsaccades occurring within a given time bin. To test the statistical significance of congruent and incongruent microsaccade proportions, we used the binomial test in each of the 50ms time windows. These analyses of microsaccade rate and direction congruence are standard analyses in the field of microsaccade research (Hafed, 2013; Tian et al., 2016). Also note that in the overt task, saccades to the images replaced post-stimulus microsaccades; it was therefore not meaningful to analyze microsaccadic modulations in the overt task.

## Results

### Both covert and overt selection behaviors are facilitated by task-irrelevant visual form images

We hypothesized that if peripheral visual forms contribute to spatial selection behaviors in an automatic and bottom-up manner, then response times associated with target detection should be influenced by the spatial congruency between target location and task-irrelevant visual form images. To test this, we first ran human subjects on two target detection tasks, one being covert and involving a manual response (Fig. 1a) and the other being overt and using a foveating eye movement response (Fig. 1b). In both tasks, the subjects had to report, as quickly as possible, the onset of a supra-threshold target stimulus that appeared at one of two possible locations centered on top of either a visual form image or its scrambled version (Methods). The subjects were informed a priori that the images behind the possible two target locations were completely irrelevant to the task, and we identified correct trials as those in which the required response (one of two buttons corresponding to each target location or accurate saccade landing at the target location) was spatially accurate; this was the majority of trials (Methods). In all cases, the target could appear after one of 3 possible delays after image onset (Fig. 1). We compared response times on correct trials when the target was congruent with the visual form image to response times when the target was incongruent with the visual form image in each delay condition.

A comparison of cumulative distributions of response times in an example subject, during the 100ms delay condition, clearly shows that response times for congruent targets were faster compared to incongruent targets in both the covert (Fig. 2a) and overt (Fig. 2d) tasks. A paired comparison of median response times across all subjects further demonstrates that response times for congruent targets were significantly faster compared to incongruent targets, in all three delay conditions (100ms, 200ms, 300ms) tested, and in both covert (Fig. 2b; Wilcoxon signed-rank test, p = 0.0009 in 100ms, p = 0.0009 in 200ms, p = 0.0009 in 300ms) and overt (Fig. 2e; Wilcoxon signed-rank test, p = 0.0019 in 100ms, p = 0.027 in 200ms, p = 0.011 in 300ms) tasks. This facilitation of response times by the visual form was uniformly present across all delays in the covert task (Fig. 2c), and it was the strongest for the 100ms delay condition in the overt task (Fig. 2f). These findings demonstrate that peripheral visual forms, even when task-irrelevant, bias spatial selection and facilitate target detection as early as 100ms from image onset, in both covert and overt spatial selection behaviors.

### Visual forms capture selection even when incongruent with task requirements

In its complementary form, a spatial selection bias by visual forms could also degrade performance and result in more error trials when visual forms are spatially incongruent with target locations. We tested for this by comparing the proportion of errors in target congruent trials with the proportion of errors in target incongruent trials, in both the covert (Fig. 3a) and overt (Fig. 3b) tasks. On average, our subjects made more errors when the targets were incongruent with visual forms compared to when they were congruent with visual forms, and this occurred in both the covert (Fig. 3a) and overt (Fig. 3b) versions of the task. A paired comparison across subjects revealed a consistent pattern of more errors for incongruent targets in the 200ms (Wilcoxon signed-rank test; covert task, p = 0.03; overt task, p = 0.01) and 300ms (Wilcoxon signed-rank test; covert task, p = 0.09; overt task, p = 0.06) delay conditions, in both the covert (Fig. 3a; middle and right panels) and overt (Fig. 3b; middle and right panels) tasks. Interestingly, this effect of visual forms on errors for incongruent targets was weaker and less consistent across subjects in the 100ms delay condition (Wilcoxon signed-rank test; covert task, p = 0.43; overt task, p = 0.22), and in both covert (Fig. 3a; left panel) and overt (Fig. 3b; left panel) tasks. These findings provide a complementary demonstration that peripheral visual forms bias spatial selection and produce more errors for incongruent targets in a time-specific manner (Fig. 2).

### Microsaccades during fixation reflect capture by peripheral, task-irrelevant visual forms

The behavioral effects of peripheral visual forms, particularly on response times (Fig. 2), in both covert and overt tasks resemble the well-known effects of spatial cues on behavioral performance in attention tasks (Posner, 1980). Since microsaccades provide a sensitive assay of effects related to attention (Hafed and Clark, 2002; Engbert and Kliegl, 2003), we therefore tested whether peripheral visual forms also bias microsaccades before target presentation. We analyzed the incidence of microsaccades before target presentation either towards (congruent) or opposite (incongruent) the suddenly appearing visual form image in the covert version of the task (Methods). We first computed a microsaccade rate independently for movements that were either congruent with the visual form image or incongruent with it (Fig. 4a). Immediately after image onset, microsaccade rate for both congruent and incongruent movements decreased reflexively, consistent with previous reports of microsaccadic inhibition (Engbert and Kliegl, 2003; Rolfs et al., 2008; Tian et al., 2016; Buonocore et al., 2017). However, subsequent microsaccades, which likely benefit from frontal cortical drive (Peel et al., 2016), occurred earlier if they were congruent with a visual form than if they were incongruent (Fig. 4a). Importantly, this meant that the proportion of microsaccades in the congruent direction was higher than in the incongruent direction during the interval following inhibition, suggesting a spatial direction bias towards visual form images. This spatial bias was statistically different from chance in each of the 50ms time bins from 122.5ms to 142.5ms (Fig. 4b; binomial test, p < 0.05). These findings demonstrate that peripheral visual forms, even when task irrelevant, bias microsaccades and effectively act as spatial cues for selection behaviors.

### Non-face visual forms strongly influence response times and bias microsaccades

The known influence of face stimuli on saccadic eye movements (Bindemann et al., 2007; Xu-Wilson et al., 2009; Morand et al., 2010; Devue et al., 2012; Boucart et al., 2016; Kauffmann et al., 2019; Buonocore et al., 2020) raises a potential question on our results so far: namely, whether our findings of visual form effects on response times, microsaccade biases, and target selection errors are largely restricted to trials with face images. To test this, we excluded trials with face images and re-analyzed all of our data with only non-face images in both covert and overt tasks. We found that response times were faster for targets congruent with non-face visual forms compared to incongruent targets in both covert (Fig. 5a; Wilcoxon signed-rank test, p = 0.002 in 100ms, p = 0.002 in 200ms, p = 0.002 in 300ms) and overt tasks (Fig. 5b; Wilcoxon signed-rank test, p = 0.01 in 100ms, p = 0.09 in 200ms, p = 0.05 in 300ms). Importantly, we also observed a spatial direction bias in microsaccades before the target presentation towards non-face visual forms in each of the 50ms bins from 112.5ms to 147.5ms (Fig. 5c; binomial test, p < 0.05). These results show that non-face visual forms strongly influence spatial selection to facilitate target detection in both covert and overt behaviors.

Additionally, we evaluated the complementary effect of non-face visual forms on errors when they were incongruent with the targets. Surprisingly, we found the effect of incongruent visual forms on errors to be weak and inconsistent across three delay conditions in both covert (Wilcoxon signed-rank test, p = 0.74 in 100ms, p = 0.21 in 200ms, p = 0.12 in 300ms) and overt tasks (Wilcoxon signed-rank test, p = 0.68 in 100ms, p = 0.007 in 200ms, p = 0.12 in 300ms). These findings suggest that non-face visual forms bias spatial selection only to an extent where it can facilitate spatially congruent target detection but not necessarily degrade spatially incongruent target detection.

### Overt selection behavior is also facilitated by peripheral visual forms in monkeys

Since monkeys are an important animal model for investigating the neural mechanisms of spatial selection behaviors (Schall and Thompson, 1999; Reynolds and Chelazzi, 2004; Krauzlis et al., 2014; Basso and May, 2017), we next asked whether peripheral visual forms can have similar effects in these animals as in our human subjects. We used the same overt task design as in humans (see Methods), and we analyzed the monkeys’ saccades. We confirmed that all three monkeys (A, F, M) performed the task correctly (% correct performance: 90.2% ± 2.1% s. d., 92.8% ± 4% s. d., and 84% ± 2.9% s. d. for monkeys A, F, and M, respectively), and we also confirmed that individual monkeys’ performance was significantly greater than chance in each of the 17 sessions collected across three monkeys (bootstrap test, p < 0.001). Following the same reasoning as in the human experiments, we compared saccadic response times on target congruent and target incongruent trials in each delay condition and monkey (Fig. 6). The results revealed two features that were consistent across all monkeys, and that were also consistent with our observations in humans when considering that monkey response times are generally faster than human response times. First, faster response times to congruent targets were limited to the early saccade responses as evident in the comparison between target congruent and target incongruent trials in the 100ms delay condition (first column of panels in Fig. 6). Second, this facilitation of early saccade responses by visual forms was very weak in the 300ms delay condition (third column of panels in Fig. 6) compared to the 100ms delay condition.

We quantified this differential effect of visual forms on early and late saccade response times by splitting the response time distributions into 7 quantiles, such that the first quantile occupied the express-saccade part of the cumulative distributions in all conditions and monkeys. Express saccades represent a population of early saccades with very short latency, which appear to be distinct from the overall response time distribution (Fischer and Boch, 1983). Thus, in the cumulative distributions of response times (e.g. Fig. 6), the express-saccade part of response time distributions appears as an early distribution of trials before a plateau is reached in cumulative response time (i.e. an early tail in the global cumulative distribution). Paired comparisons of median response times for target congruent and target incongruent trials in the 1^st^ quantile showed that saccadic responses were significantly faster for congruent targets in the 100ms and 200ms delay conditions (Fig. 7a; Wilcoxon signed-rank test, p = 0.001 in 100ms, p = 0.003 in 200ms) but not in the 300ms delay condition (Fig. 7a; Wilcoxon signed-rank test, p = 0.129), consistent with the observations from Fig. 6. The effect on all ranges (quantiles) of saccade response times in the three delay conditions is also shown in Fig. 7b. As can be seen, there was a facilitatory effect of peripheral visual forms on saccade response times, but this was limited to the early saccadic responses and fell off abruptly for the 300ms delay condition after the 1^st^ quantile. The fall off was milder for the 100ms and 200ms delay conditions (Fig. 7b). These findings demonstrate that task-irrelevant visual forms facilitate early saccade responses in monkeys, and that this facilitation is the strongest in the first 200ms of visual form processing, consistent with our earlier results in humans (Fig. 2f).

**Figure 7.**
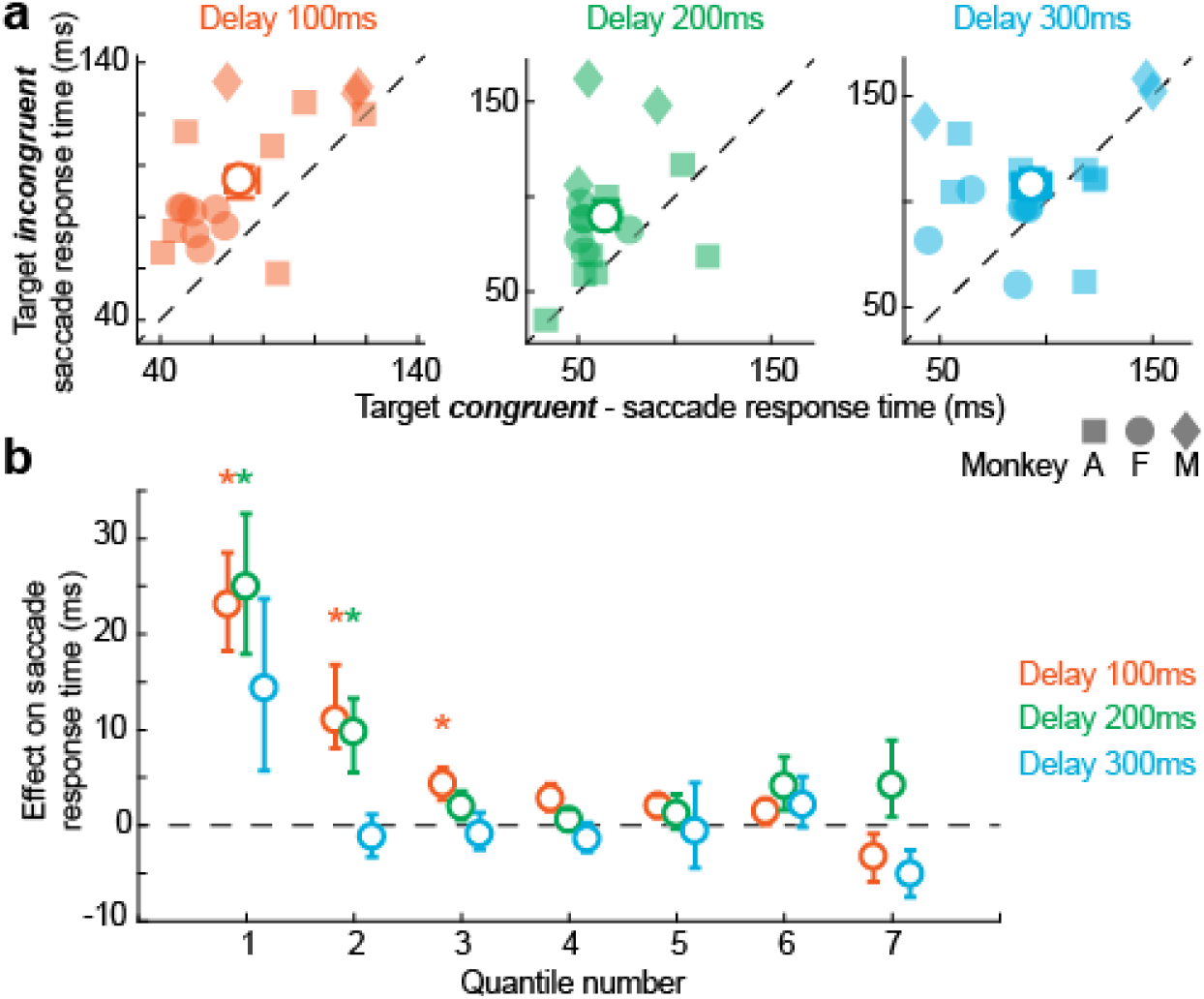
Visual form facilitation of early saccade responses in the overt selection task in monkeys. **(a)** Response times in each monkey were split into seven quantiles (see Methods). Each panel shows a paired comparison of median response times in the first quantile between target congruent and target incongruent trials for a fixed delay of 100ms (red), 200ms (green), or 300ms (blue) across all sessions (n = 17; filled symbols) from three monkeys. The colored circle with errors bars in each panel represents mean and standard deviation across sessions. All monkeys showed faster response times for target congruent trials than target incongruent trials, consistent with our human results (Fig. 2). **(b)** The effect of visual forms on saccade response times was quantified in each response time quantile as a difference in median response time between incongruent and congruent trials in the corresponding quantile. The mean effect on response times across sessions (colored circles) is shown separately for all seven quantiles and the three fixed delays (see legend). Error bars indicate standard deviation across sessions. * indicates significance using the Wilcoxon signed-rank test in the corresponding delay and quantile (p < 0.05). Trials with fast response times were consistently associated with a facilitatory effect of visual forms on overt spatial selection behavior in monkeys.

### Visual forms capture more saccade errors in monkeys when incongruent with task requirements

Finally, in humans, the visual form facilitation of response times for target congruent trials was accompanied by the complimentary effect of more errors for targets that were incongruent with visual forms images (Fig. 3). We tested whether peripheral visual forms had this complementary effect on errors in monkeys as well. We found that monkeys indeed made significantly more errors when the saccade targets were incongruent with the visual forms compared to when the targets were congruent with the visual forms (Fig. 8). Interestingly, this effect on errors was strong and significant in the 100ms (Wilcoxon signed-rank test, p = 0.0006) and 200ms (Wilcoxon signed-rank test, p = 0.02) delay conditions but relatively weak and insignificant in the 300ms delay condition (Wilcoxon signed-rank test, p = 0.45). This stronger effect on errors in the early delay conditions is consistent with the similar results from the saccade response times (Fig. 7). These findings demonstrate that peripheral visual forms capture more error saccades in well-trained monkeys, and that this effect on errors is the strongest in the first 200ms of visual form processing.

**Figure 8.**
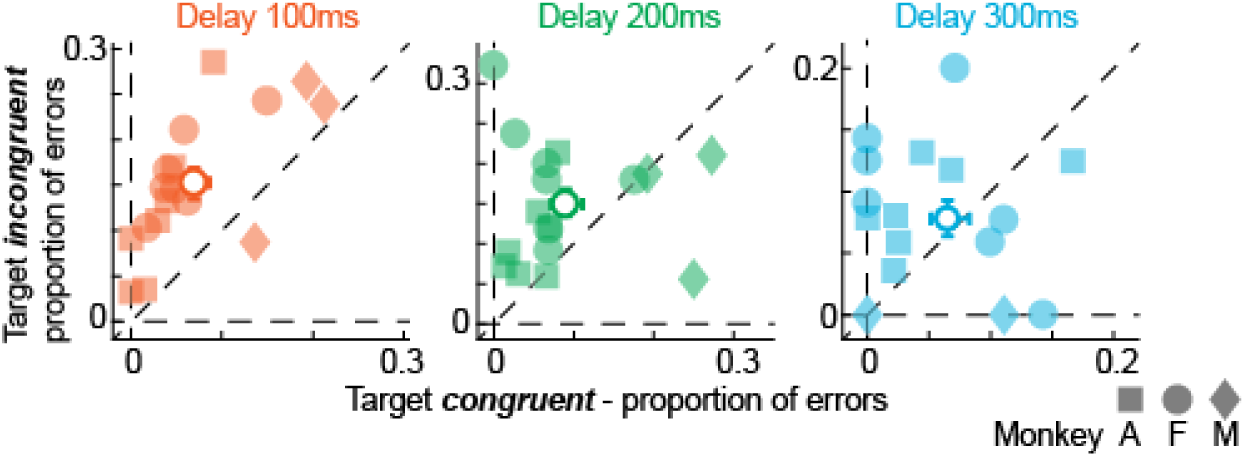
Visual form influence on errors in the overt spatial selection task in monkeys. Each panel shows a paired comparison of proportion of saccade response errors between target congruent and target incongruent trials for a fixed delay of 100ms (red), 200ms (green), or 300ms (blue) across all sessions (n = 17; filled symbols) from three monkeys (see legend). The colored circle with errors bars in each panel represents mean and standard deviation across sessions. All monkeys had more errors in the target incongruent condition than in the target congruent condition, consistent with our human observations (Fig. 3).

Importantly, we also confirmed in monkeys that non-face visual forms still strongly influenced response times in the 1^st^ quantile (Wilcoxon signed-rank test, p = 0.008 in 100ms, p = 0.006 in 200ms, p = 0.18 in 300ms) and errors (Wilcoxon signed-rank test, p = 0.0004 in 100ms, p = 0.03 in 200ms, p = 0.39 in 300ms) in the first 200ms of visual form processing, as demonstrated in our main findings (Figs. 7a, 8).

## Discussion

We investigated whether peripheral visual forms contribute to covert and overt spatial selection behaviors using a visual detection task in which visual forms were completely irrelevant. In humans, we found that visual forms facilitate the detection of spatially congruent targets with faster response times in both covert (Fig. 2b) and overt (Fig. 2e) tasks, and that this facilitation is evident in the first 100ms of visual form processing. Importantly, visual forms incongruent with targets resulted in more errors in both covert (Fig. 3a) and overt (Fig. 3b) tasks, and this effect on errors was most pronounced after 200ms of visual form processing. In addition, microsaccades before target presentation (but after visual form image onset) were biased towards visual forms in the covert task (Fig. 4b). Our results in monkeys revealed a similar pattern of visual form effects seen in humans with two notable differences. First, visual form facilitation of response times was specific to early saccadic responses (Fig. 7b), likely because monkey saccadic response times are faster than those of humans. Second, the visual form effects on response times and errors were limited to the first 200ms of visual form processing (Figs. 7, 8). Overall, these findings demonstrate that peripheral visual forms contribute to covert and overt spatial selection in ways that resemble the effects of spatial cues on orienting behaviors (Posner, 1980).

### Low-level visual factors cannot explain visual form influences on response times

Low-level visual factors related to luminance, spatial frequency content, and target contrast modulate neuronal activity in visual and sensorimotor regions of the brain (Ohayon et al., 2012; Chen and Hafed, 2018; Chen et al., 2018; Vinke and Ling, 2020), and therefore may influence behavioral responses. For this reason, we took several measures in the design of the image and target stimuli to minimize the contribution of low-level visual factors to our findings, particularly on response times. First, we equalized all visual form images and their corresponding 50% phase-scrambled images iteratively for luminance distributions and the Fourier spectra (see Methods). Second, we chose the target to be of the highest contrast and adjusted the size so that target detection and hence the perceived contrast of the target, was supra-threshold for both visual form and phase-scrambled image backgrounds (see Methods). Thus, we suggest that low-level visual factors were unlikely to have influenced our results showing visual form facilitation of response times.

### High-level cognitive factors cannot explain visual form influence on response times

Cognitive factors related to behavioral relevance, novelty, and reward also modulate neuronal activity in visuomotor brain regions, such as the superior colliculus, and hence can shape orienting behaviors (Basso and Wurtz, 1997; Ikeda and Hikosaka, 2003; Boehnke et al., 2011; Herman and Krauzlis, 2017). These factors are again unlikely to have influenced our findings for the following reasons. First, we made the visual form and the phase-scrambled images completely irrelevant to behavior in both covert and overt tasks, and in both humans and monkeys. Second, the same subjects participated in both covert and overt tasks that used the same images (see Methods). In addition, all monkeys were trained with the same images in at least 7 training sessions before the experimental sessions. Third, none of the images were associated with reward, in humans and monkeys, as they were irrelevant to the performance in the task. Thus, cognitive factors related to behavioral relevance, novelty, and reward were unlikely to have influenced our findings showing visual form facilitation of response times.

### Face stimuli alone cannot account for visual form influence on response times

Faces are of ecological value, and the influence of faces on goal-directed and free-viewing saccade behaviors is well documented (Bindemann et al., 2007; Xu-Wilson et al., 2009; Morand et al., 2010; Devue et al., 2012; Boucart et al., 2016; Kauffmann et al., 2019; Buonocore et al., 2020). Importantly, there is growing evidence that faces are rapidly processed through a network of subcortical structures including the superior colliculus (Johnson, 2005; Nguyen et al., 2016; Le et al., 2020), which also plays a crucial role in spatial selection (McPeek and Keller, 2004; Lovejoy and Krauzlis, 2009). To confirm that face images alone did not disproportionately contribute to our results, we repeated all of our analyses of covert and overt tasks in humans by excluding trials with face images. Results showed that response times were equally strongly affected by non-face visual forms alone in both covert (Fig. 5a) and overt tasks (Fig. 5b). Importantly, we also observed significant biases in microsaccades to non-face visual forms (Fig. 5c). Additionally, we also confirmed in monkeys that non-face visual forms strongly influenced the response times. These control analyses show that face stimuli alone cannot account for visual form influence on response times in both covert and overt tasks, and most importantly, demonstrate that all visual forms can influence spatial selection.

### Comparison of visual form effects in humans and monkeys

A comparison of visual form effects in humans and monkeys during the overt task revealed interesting species differences. For example, visual form effects on response times in monkeys were confined to the earliest saccades (Fig. 7b) unlike in humans where a similar analysis revealed visual form effects across all quantiles in all delay conditions (Fig. 2b). Similarly, visual form effects on errors in monkeys were more pronounced in the early delay conditions (100ms and 200ms; see Fig. 8) with a weaker effect in the late 300ms delay condition, unlike in humans where this pattern was almost reversed; effects on errors were the weakest in the 100ms condition (Fig. 3b). We suggest that the predominance of visual form effects on earliest responses and delay conditions in monkeys may be related to their behavioral training. Specifically, these were highly trained animals with short saccadic response times in general. With the longer delay periods (e.g. 200ms and 300ms), these delay periods were often much longer than the actual saccadic reaction times that would have been elicited to the visual form images themselves (e.g. see Fig. 7a). The long delay periods therefore required actively suppressing saccades to properly receive rewards in the task, which eliminated the visual form effects that still occurred automatically with shorter latencies. Indeed, the monkeys’ final reaction times on successful trials were much shorter than those of the human subjects in the same task (compare Figs. 2 and 7).

### Visual-form based selection differs from object-based attention

Space-based or spatial attention refers to behavioral benefits conferred by spatial cues exclusively at the cued location (Carrasco, 2011). In object-based attention, the cueing benefits extend to all spatial locations occupied by the object at the cued location (Duncan, 1984; Egly et al., 1994; Abrams and Law, 2000). Our demonstration of spatial selection based on visual forms is different from object-based attention because there were no explicit spatial cues in our task, and, most importantly, the visual forms were irrelevant in our task. However, it is very likely that both object-based and visual-form based selection mechanisms involve common visual processes related to segmentation and perceptual grouping (Driver et al., 2001; Baldauf and Desimone, 2014), and may operate outside of the modulation of sensory processing mechanisms associated with spatial attention (Shomstein and Yantis, 2002; Reynolds and Chelazzi, 2004; Chou et al., 2014); but also see (Roelfsema et al., 1998).

### Neural circuits representing visual-form based spatial maps for orienting

The influence of peripheral visual forms on target detection as early as 100ms suggests a neural circuit that rapidly links visual form processing with spatially maps in sensorimotor structures, such as the superior colliculus (Robinson, 1972; Chen et al., 2019). Recent evidence in a new region of the primate temporal cortex shows rapid object-selectivity and detection-related signals that were causally dependent on midbrain superior colliculus activity (Bogadhi et al., 2019b). Based on this evidence, we suspect that superior colliculus neurons might signal peripheral visual forms and bias spatial selection. Recent findings in monkeys and mice further demonstrate the visual capabilities of superior colliculus neurons in representing visual statistics and properties of the natural environment that are innately relevant to our behaviors (Hafed and Chen, 2016; Chen et al., 2018; Lee et al., 2020).

Of course, visual form recognition is also accomplished in the primate inferotemporal (IT) cortex through feed-forward visual cortical circuits. This can possibly influence sensorimotor structures such as the superior colliculus for spatial selection through direct projections (Cerkevich et al., 2014). However, the time course of visual form recognition in the traditional IT regions is not entirely consistent with our results showing rapid visual form facilitation (Kreiman et al., 2006; Tsao et al., 2006). Therefore, we suggest that a circuit linking superior colliculus with the temporal cortex, possibly through pulvinar or amygdala, may be at play in linking rapid visual form recognition with spatial selection (Harting et al., 1991; Boussaoud et al., 1992; Hadj-Bouziane et al., 2012; Rafal et al., 2015; Soares et al., 2017). Future studies investigating the subcortical and cortical contributions to visual form recognition, particularly in the periphery, will identify the candidate circuit mediating the visual form influence on spatial selection.

## Acknowledgements

We were funded by the Werner Reichardt Centre for Integrative Neuroscience, an excellence cluster funded by the Deutsche Forschungsgemeinschaft (DFG; EXC 307). We were also funded by the Hertie Institute for Clinical Brain Research, and DFG project BO5681/1-1.

